# Simulating the Impact of Tumor Mechanical Forces on Glymphatic Networks in the Brain Parenchyma

**DOI:** 10.1101/2024.05.18.594808

**Authors:** Saeed Siri, Alice Burchett, Meenal Datta

## Abstract

**Background:** The brain glymphatic system is currently being explored in the context of many neurological disorders and diseases, including traumatic brain injury, Alzheimer’s disease, and ischemic stroke. However, little is known about the impact of brain tumors on glymphatic function. Mechanical forces generated during tumor development and growth may be responsible for compromised glymphatic transport pathways, reducing waste clearance and cerebrospinal fluid (CSF) transport in the brain parenchyma. One such force is solid stress, i.e., growth-induced forces from cell hyperproliferation and excess matrix deposition. Because there are no prior studies assessing the impact of tumor-derived solid stress on glymphatic system structure and performance in the brain parenchyma, this study serves to fill an important gap in the field.

**Methods:** We adapted a previously developed Electrical Analog Model using MATLAB Simulink for glymphatic transport coupled with Finite Element Analysis for tumor mechanical stresses and strains in COMSOL. This allowed simulation of the impact of tumor mechanical force generation on fluid transport within brain parenchymal glymphatic units – which include paravascular spaces, astrocytic networks, interstitial spaces, and capillary basement membranes. We conducted a parametric analysis to compare the contributions of tumor size, tumor proximity, and ratio of glymphatic subunits to the stress and strain experienced by the glymphatic unit and corresponding reduction in flow rate of CSF.

**Results:** Mechanical stresses intensify with proximity to the tumor and increasing tumor size, highlighting the vulnerability of nearby glymphatic units to tumor-derived forces. Our stress and strain profiles reveal compressive deformation of these surrounding glymphatics and demonstrate that varying the relative contributions of astrocytes vs. interstitial spaces impact the resulting glymphatic structure significantly under tumor mechanical forces. Increased tumor size and proximity caused increased stress and strain across all glymphatic subunits, as does decreased astrocyte composition. Indeed, our model reveals an inverse correlation between extent of astrocyte contribution to the composition of the glymphatic unit and the resulting mechanical stress. This increased mechanical strain across the glymphatic unit decreases the venous efflux rate of CSF, dependent on the degree of strain and the specific glymphatic subunit of interest. For example, a 20% mechanical strain on capillary basement membranes does not significantly decrease venous efflux (2% decrease in flow rates), while the same magnitude of strain on astrocyte networks and interstitial spaces decreases efflux flow rates by 7% and 22%, respectively.

**Conclusion:** Our simulations reveal that solid stress from brain tumors directly reduces glymphatic fluid transport, independently from biochemical effects from cancer cells. Understanding these pathophysiological implications is crucial for developing targeted interventions aimed at restoring effective waste clearance mechanisms in the brain.

This study opens potential avenues for future experimental research in brain tumor-related glymphatic dysfunction.

## Background

The glial-associated lymphatic system, known as the glymphatic system, has received substantial attention in recent years [1-4]. The glymphatic system facilitates metabolite clearance and interstitial fluid regulation; interruption of this flow causes the buildup of toxic species and neurological dysfunction and is associated with a range of neurological diseases [5-8]. While studies to understand the significance of these networks in maintaining brain health and function are ongoing [9, 10], there is limited knowledge regarding glymphatic function in the presence of a brain tumor. In addition to biochemical effects from cancer cells, tumor-induced mechanical forces may damage the brain [11] and severely impact waste clearance in brain tumor patients [12-18]. However, the mechanisms of this effect have yet to be fully elucidated [2].

Fluid dynamics in the brain parenchyma involves flow through vascular spaces, paravascular spaces, intracellular flow through cellular networks (e.g., astrocytes), and extracellular flow in interstitial space [2]. Cerebrospinal fluid (CSF) originates in the choroid plexus, enters the brain parenchyma and travels along the periarterial space of arterioles, and ultimately drains via perivenous space of venules into the subarachnoid plexus [5]. Two parallel paths of fluid flow are hypothesized to allow flow between the periarterial and perivenous spaces. Gaps between perivascular astrocyte end-feet allow the interchange of fluid between the perivascular space and the interstitial space, where the fluid encounters various brain cell types and extracellular matrix (ECM) components [19, 20]. Water can also flow through an intracellular pathway, entering astrocytes via aquaporin-4 (AQP4) channels and moving through gap junctions between cells [5, 21]. These pathways are indirectly related, as alterations in the intracellular pathway significantly alters flow through the interstitial pathway [22]. While *in vivo* studies using injected tracers have been informative in understanding the glymphatic system, cell-scale transport and transport of smaller solutes cannot always be directly measured; thus some recent insights into the mechanisms and interactions in glymphatic flow have been found only through mathematical modelling informed by experimental observations [5, 21].

As brain tumors grow, they exert fluid (i.e., “edema) and solid (i.e., “mass effect”) stresses both intratumorally and extratumorally on the surrounding brain tissue [11, 23-25]. Brain tumor growth not only interferes with neurological function [26-31], but hinders glymphatic function as well [14] (Fig. 1). Across cancer types, tumors are known to alter interstitial fluid dynamics [32]. This is best illustrated by the edema that results from brain tumors, which is associated with worse prognosis and decreased glymphatic function [13, 18]. *In vivo* studies of a range of brain tumor types consistently demonstrate reduced rates of tracer influx and outflux, with a net lower rate of clearance and increased residual tracer [12-14, 16-18]. This may be mediated in part by decreased AQP4 expression in perivascular astrocytes and heavily disrupted vasculature associated with brain tumors [14, 24, 33, 34].

**Figure 1:**
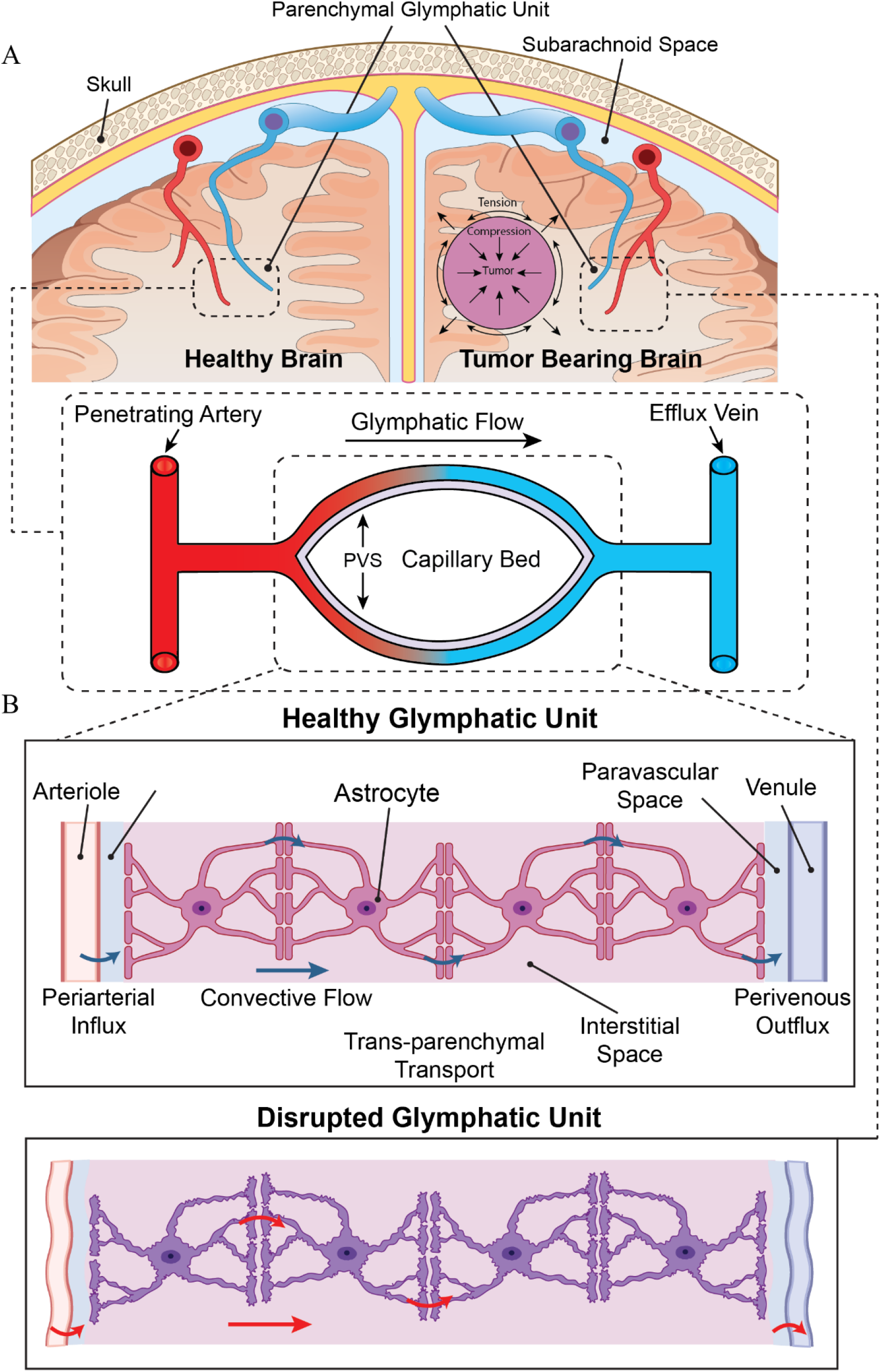
Schematic Representation of a Healthy and a Tumor-Bearing Brain, Highlighting the Glymphatic Unit and its Subcomponents. (A) The healthy brain, (left) has normal glymphatic units that clear waste products from the brain. Glymphatic flow begins along the periarterial space, proceeds through the interstitium, and exits via the perivenous space. The presence of a tumor (right) causes growth-induced solid stress; these physical forces exerted outwards on the surrounding brain can mechanically disrupt this system. Tumors likely induce biochemical alterations in the glymphatic unit as well, which is captured in our model parameters. (B) The glymphatic unit is composed of the periarterial space along the penetrating arteriole, capillary basement membrane, astrocytes, interstitial space, and perivenous space along the exiting venule. Together, these comprise pathways for trans-parenchymal transport of cerebral spinal fluid (CSF). Tumors disrupt this glymphatic unit, altering the paravascular, astrocyte, and interstitial fluid pathways.

Recent computational models have demonstrated the importance of various components of the glymphatic transport pathway [22, 35-37]. However, there is a gap in the field regarding the mechanical impact of a tumor on the brain glymphatic system, i.e., the relationship between aberrant solid stress and CSF flow in the brain. Therefore, we adapted prior models to form a comprehensive computational model to identify the most important physical parameters in glymphatic disruption with brain tumors. We found that tumor size and location determine the degree of deformation and mechanical stress experienced by the glymphatic system. We also found that increased astrocyte content serves to protect the glymphatic system from the detrimental effects of a tumor, while reduced astrocyte content makes it more vulnerable. Finally, glymphatic compression due to proximity to a tumor decreases CSF flow, an effect mediated by the interstitial elements and not the capillary basement membrane.

## Methods

### Finite Element Method (FEM) modeling of the glymphatic system structure

The governing equations for FEM structural analysis involve equilibrium equations and constitutive relations. The equilibrium equations represent the balance of forces within the brain parenchyma. They can be expressed as:

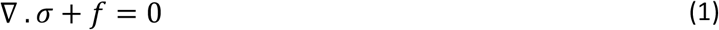

Where *σ* is the stress tensor representing internal forces and *f* is the body force per unit volume. Constitutive relations describe the relationship between stress and strain in the material. For linear elasticity, Hooke’s Law is commonly used:

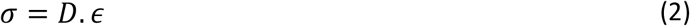

Where *ϵ* is the strain tensor representing deformations, and *D* is the elasticity tensor (material property). Strain-Displacement Relation defines the relationship between strain and displacement. For small deformations:

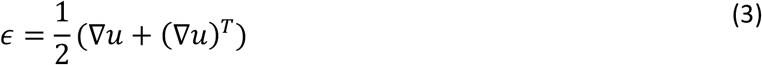

Where *u* is the displacement vector. When these equations are discretized using FEM, the continuous domain of the brain parenchyma is divided into smaller, simpler elements. The displacement field within each element is approximated using shape functions, and the equilibrium and constitutive equations are formulated for each element. Together, these equations generate a system of linear equations that we solve numerically to determine the displacement and stress distribution within the brain parenchyma under given loading and boundary conditions.

A 2D geometry (Fig. 2B) including brain astrocytes, IS, BM, PVS, and brain tumors was constructed for FEM modeling using COMSOL Multiphysics. The FEM model allowed for the simulation of a growing nodular brain tumor (with pushing and not replacing behavior [11]), with the initial tumor radius of r_0_ = 1 mm, incorporating the geometry and mechanical properties of all glymphatic unit subcomponents. Within the FEM framework, parametric analyses were conducted to evaluate the contributions of astrocytes and interstitial spaces, and their mechanical and geometrical characteristics, as well as tumor size and proximity to the representative parenchymal glymphatic unit. Specifically, variations in astrocyte and IS composition (including ECM contribution alterations) were simulated to understand their impact on glymphatic flow. The parametric analysis for the glymphatic unit mechanical structure was conducted for 10 different scenarios (Figs. 3 and 4): 10% astrocyte - 90% IS to 90% astrocyte - 10% IS (by %10 percent increments). We assumed linear elastic mechanical behavior for this initial model development and implementation, given the current lack of definitive single cell resolution experimental viscosity and relaxation time data for the model components [38]. The Young’s moduli of astrocytes, interstitial space, and basement membrane were considered E= 850 Pa, 300 Pa, and 400 Pa respectively [39, 40].

**Figure 2:**
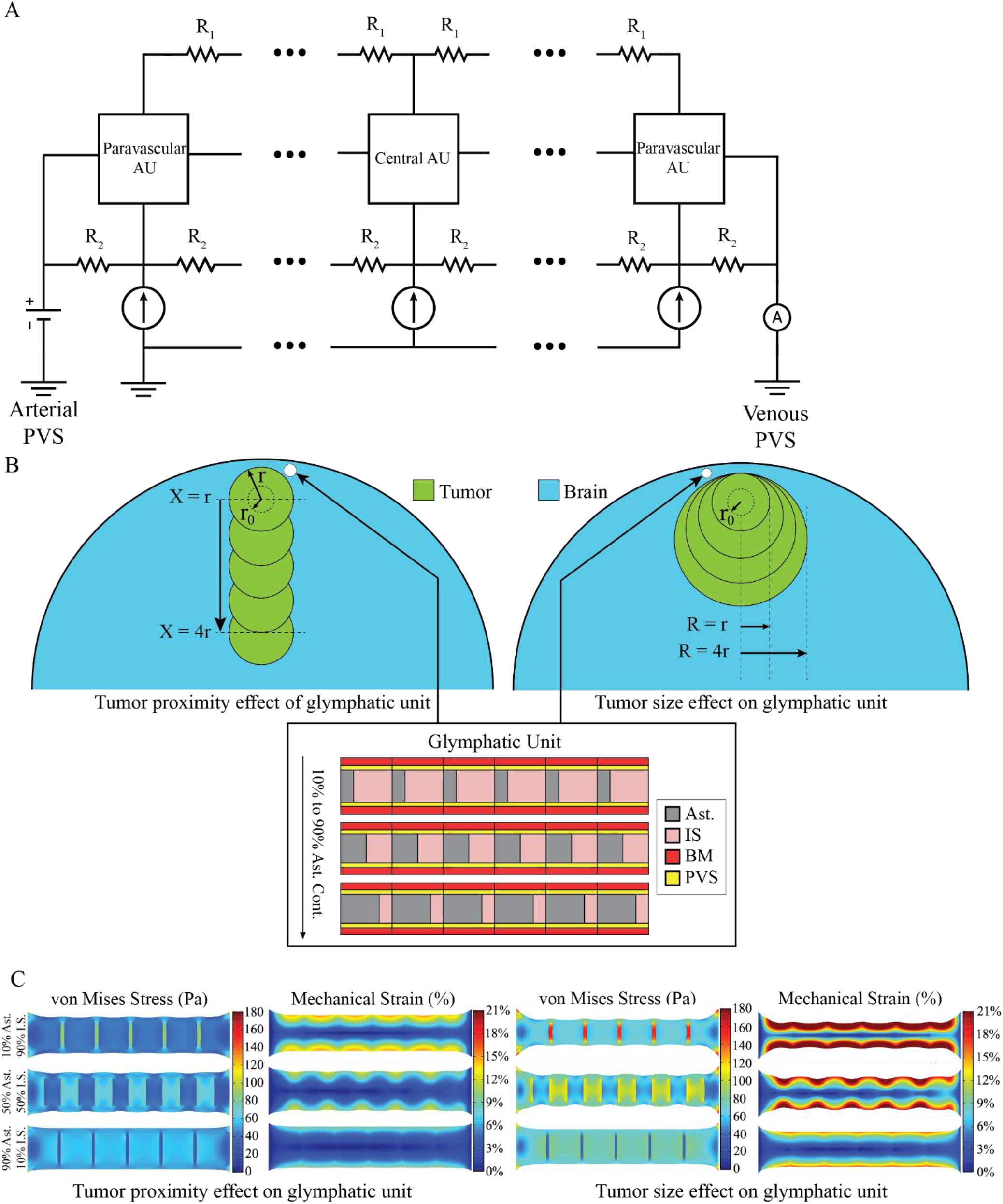
Schematic of the Computational Model of Fluid Flow and Solid Stress in the Brain Tumor-Adjacent Glymphatic Unit. (A) The implemented electrical analogue model of cerebral water transport between arterial and venous paravascular spaces including two paravascluar units and four central units. R_1_ represents the gap junction resistance and R_2_ represents basement membrane resistance. The potential difference across the circuit represents driving fluid pressure difference between the perivenous and periarterial space, and the current of the circuit represents fluid flow. Current flows from the paravascular space across paravascular astrocyte units (AUs), through four sequential central AUs, and through a second paravascular AU before exiting. Redrawn from ref. [22] (B) 2D geometric representations of the brain and tumors tailored for parametric analysis using Finite Element Method (FEM) modeling in COMSOL Multiphysics. To study the effect of tumor proximity, simulations are performed from far (X=4r) from the glymphatic unit to adjacent (X=0) to the glymphatic unit. To study the effect of tumor size, simulations are performed from tumor sizes R=r to R=4r for tumors adjacent to the glymphatic unit. (Ast. – astrocytes, IS – interstitial space, BM – capillary basement membrane, PVS – paravascular space). (C) Representative finite element simulations for the parametric analysis showing the mechanical stress and strain map with varying contributions of the three major components of the glymphatic system. Tumor Proximity (left) and Tumor Size (right).

**Figure 3:**
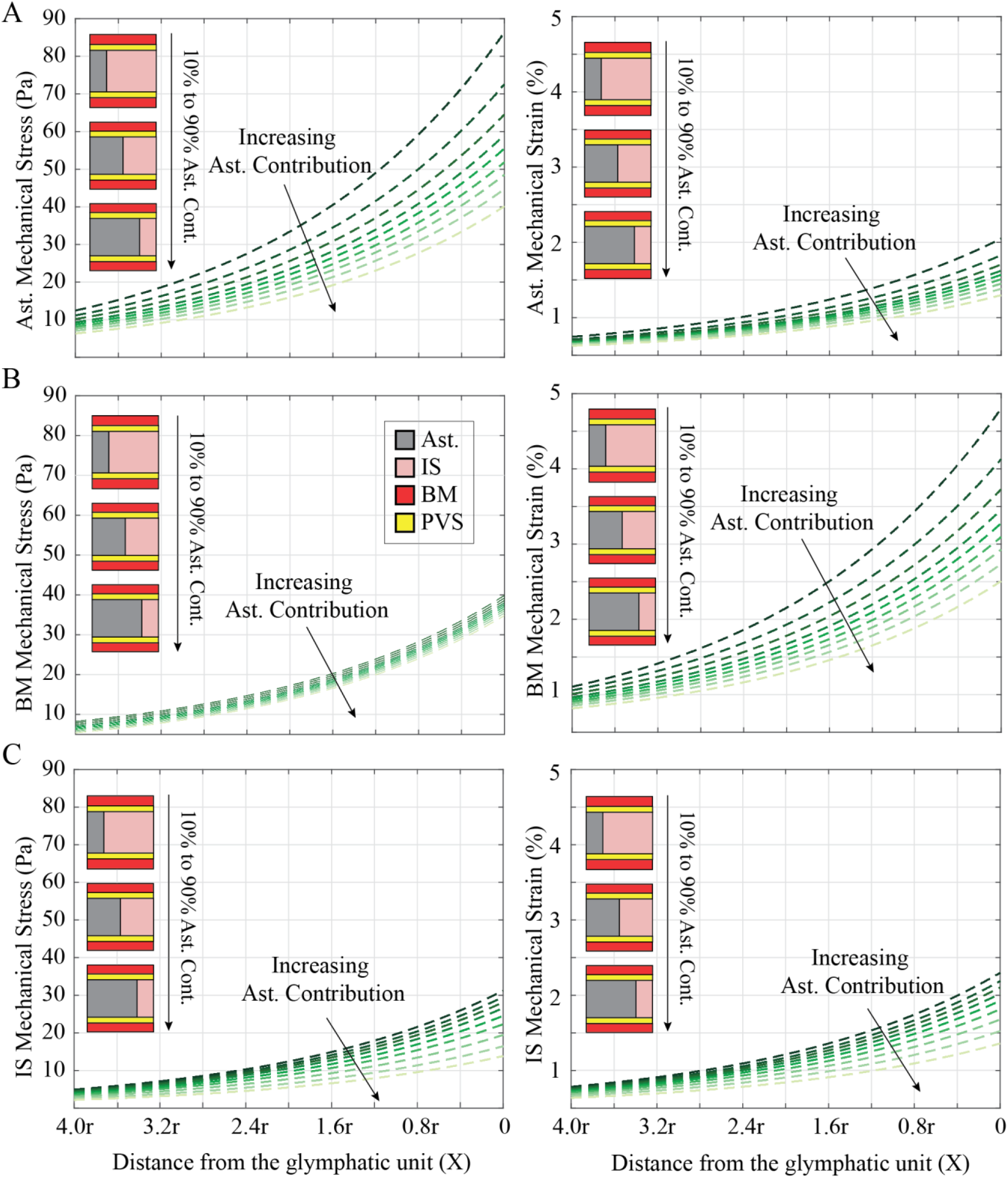
Brain Tumor Proximity Influences Stress and Strain of Glymphatic Subunits. The von Mises mechanical stress experienced by the glymphatic subunits as a function of the distance of the tumor from the glymphatic unit. The simulations are performed from far (X=4r) from the glymphatic unit to adjacent (X=0) to the glymphatic unit. Von Mises stress and mechanical strain in the (A) astrocytes (Ast.), (B) basement membrane (BM), and (C) interstitial space (IS) as a function of tumor proximity. Each condition includes 10 simulations ranging from 10% Ast.:90% IS contribution to 90% Ast.:10% IS contribution.

**Figure 4:**
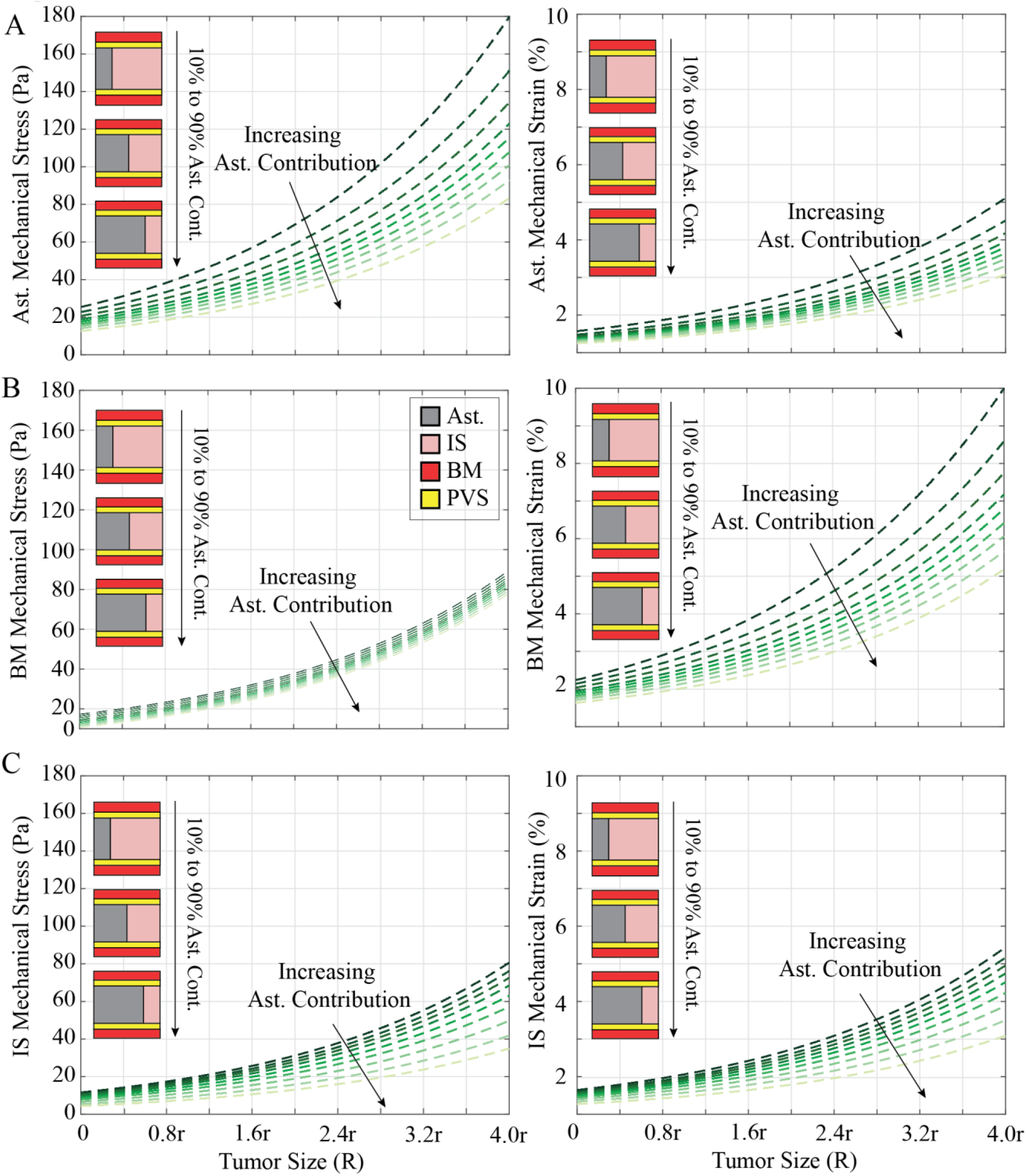
Brain Tumor Size Influences Stress and Strain of Glymphatic Subunits. The von Mises mechanical stress experienced by the glymphatic unit as a function of the tumor size adjacent to the glymphatic unit. The simulations are performed for tumor sizes R=r to R=4r. Von Mises stress and mechanical strain in the (A) astrocytes (Ast.), (B) basement membrane (B.M.), and (C) interstitial space (I.S.) as a function of tumor size. Each condition includes 10 simulations ranging from 10% Ast.:90% IS contribution to 90% Ast.:10% IS contribution.

### Parametric analysis of the tumor location and size

We calculated the average mechanical stress and strain exerted on the glymphatic system by assuming the glymphatic unit’s placement “atop and adjacent” to the tumor (Fig. 2B). Our modeling considered two key factors: (i) the tumor location, ranging from immediately beside the glymphatic unit to a distance of 4r, where r is the tumor radius; and (ii) the tumor radius, varying from r = 10 mm to 4r = 40 mm. We also calculated the mechanical stress and strain map for varying structural contributions of the three major components of the glymphatic system to assess the effects of tumor proximity and tumor size on glymphatic system (Fig. 2C).

### Electrical analog model of glymphatic fluid transport

We built upon a previously published glymphatic network model by Asgari et al. [22] and applied relevant geometric and physical properties associated with the presence of brain tumors. This electrical analog model considers a representative parenchymal glymphatic unit as blood capillary basement membranes (BM) and paravascular spaces (PVS), surrounding a network of astrocytes, and interstitial spaces (IS) (Fig. 2A). We implemented this model in MATLAB Simulink to analyze the impact of brain tumors on glymphatic transport and explore how alterations in the astrocyte network contribute to compromised fluid flow in the brain parenchyma (Figs. 3 and 4).

### Derivation of venous side flow rate based of the variations of mechanical strain

By employing an integrated approach, we coupled Finite Element Method (FEM) modeling with the predeveloped glymphatic system lumped parameter model, using the COMSOL’s MATLAB Live Link package. That enabled us to examine the effects of the mechanical strain on the geometry of the glymphatic system’s subcomponents. These changes, in turn, impact the electrical resistance within the system, leading to subsequent adjustments in the venous side flow rate. Several key parameters govern the behavior of the glymphatic system, each with its associated equations and nominal values. These parameters include the distance between arteriole-venule pairs, capillary diameter, astrocyte soma diameter, volume, surface area, and various other properties that influence the astrocyte’s role in modulating fluid flow and solute transport, and also IS thickness, capillary BM thickness and hydraulic permeability. (See Table 2 in ref. [22] for details.)

## Results

### Tumor proximity and size to the glymphatic unit significantly elevates von Mises stress and mechanical strain across all components

Fig. 2C depicts the von Mises stress and mechanical strain profiles in three distinct scenarios characterized, arising from varying the proportion of astrocytes and the IS. In the first scenario, where astrocytes contribute minimally in the glymphatic unit structure and IS assumes a more substantial role, the glymphatic unit structure exhibits the highest strain profiles, primarily attributed to IS (10% averaged mechanical strain through the whole structure). In the second scenario, featuring a more balanced contribution between astrocytes and IS in the glymphatic unit structure, the strain profiles are distributed uniformly across the structure, yet with a predominant concentration in the IS. Conversely, in the third scenario, where astrocytes play a more significant role with the lowest contribution from IS, the results demonstrate minimal averaged mechanical strains to the glymphatic unit compared to the other two scenarios. Intriguingly, the mechanical stress on astrocytes peaks when their contribution is at its lowest, and it is at its lowest when their contribution is most substantial.

In the astrocyte subunit, when tumors are close to the unit and astrocytes contribute minimally (10%), astrocytes experience the highest von Mises mechanical stress, reaching approximately ∼85 Pa (Fig. 3A). Conversely, an increased astrocyte content (90%) mitigates the effect of the tumor, reducing von Mises stress to ∼40 Pa across the astrocytes. In the astrocyte subunit, the maximum stress value doubles from ∼85 Pa to ∼170 Pa for adjacent tumors with sizes R = r and R = 4r, respectively (Fig. 4A). Again, increasing astrocyte contribution reduces stress and strain across all subunits and mitigates the effect of increasing tumor size. Increasing tumor size from R = r to R = 4r in the case of 90% astrocytes results in a ∼65 Pa increase in astrocyte stress, while decreasing astrocyte content from 90% to 10% results in a ∼100 Pa increase in astrocyte stress at a tumor size of R = 4r. Thus, astrocyte composition is as or more important than tumor size and proximity in determining astrocyte mechanical stress.

### The influence of tumor proximity and size on the glymphatic unit mechanical stress and strain varies across its subcomponents

Within the astrocyte subunit (Fig. 3A), von Mises stress spans from approximately 5 to 40 Pa with the highest astrocyte contribution, and from 12 to 87 Pa with the lowest astrocyte contributions. The associated mechanical strain ranges from approximately 0.2% to 1.6% for the highest astrocyte contribution and about 0.3% to 2% for the lowest astrocyte contributions. When the tumor is far from the glymphatic unit (4r), the basement membrane (BM) (Fig. 3B) experiences von Mises stress ranging from approximately 0 Pa to 7 Pa with 90% and 10% astrocyte contribution, respectively. When the tumor is adjacent to the glymphatic unit, the BM stress ranges from approximately 35 Pa to 40 Pa with 90% and 10% astrocyte contribution, respectively. Tumor proximity is clearly the most important determining factor in BM stress, with changing astrocyte content only altering stress values by 5-7 Pa, regardless of proximity. However, astrocyte’s contribution plays a more significant role in BM strain. When the tumor is adjacent, decreasing astrocyte content from 90% to 10% results in an increase in strain from 2.5% to 4.75%, while increasing proximity increases strain by 2-3.75% for 10-90% astrocyte contribution. In the interstitial space (Fig. 3C), von Mises stress ranges approximately from 1 to 13 Pa with the highest astrocyte contribution and varies from about 3 to 31 Pa with the lowest astrocyte contributions. The mechanical strain in this space ranges from approximately 0.2% to 1.8% for the highest astrocyte contribution and about 0.5% to 2.7% for the lowest astrocyte contributions.

In the parametric analysis of tumor size, for the largest tumor adjacent to the glymphatic unit in the astrocytes (Fig. 4A), von Mises stress spans from approximately 11 to 83 Pa with the highest astrocyte contribution, while it varies from about 26 to 180 Pa with the lowest astrocyte contributions. The associated mechanical strain ranges from approximately 0.5% to 3.1% for the highest astrocyte contribution and about 1% to 5.1% for the lowest astrocyte contributions. The BM (Fig 4B) exhibits von Mises stress ranging approximately from 0 to 85 Pa with the highest astrocyte contribution, and it varies from about 18 to 100 Pa with the lowest astrocyte contributions. Correspondingly, the mechanical strain ranges from 1.2% to 5.2% for the highest astrocyte contribution and approximately 2.3% to 10% for the lowest astrocyte contributions. As with the proximity analysis, astrocyte contribution is less important than tumor size with respect to stress but becomes as important as size with respect to strain. In the interstitial space (Fig. 4C), von Mises stress ranges approximately from 2 to 38 Pa with the highest astrocyte contribution and varies from about 12 to 80 Pa with the lowest astrocyte contributions. The mechanical strain in this space ranges from approximately 0.5% to 3.1% for the highest astrocyte contribution and about 1.3% to 5.6% for the lowest astrocyte contributions.

### Mechanical compression from tumors correlates with reduced flow rate across all glymphatic unit subcomponents

We explored the functional consequences of mechanical compression on the glymphatic fluid transport using our coupled FEM and lumped parameter simulations of the solute transport in the glymphatic unit. The tumor-induced mechanical stress and strain affects the geometry of the glymphatic system and its subunits. These changes, in turn, impact the electrical resistance within the glymphatic system’s lumped parameter model, leading to subsequent adjustments in the venous side flow rate within the astrocytes, interstitial space, and capillary basement membrane. Fig. 5 depicts how different levels of mechanical strain applied to each individual subunit, ranging from 0 to 20%, impact venous side flow rate efflux, a key measure of waste clearance efficiency. Increasing mechanical strain reduces flow rate across all glymphatic unit subcomponents. The capillary BM exhibited a roughly 2% decrease in flow rate at 20% strain, while astrocytes and IS showed more substantial reductions of approximately 7% and 22%, respectively. For example, in the case of 10% astrocyte contribution, a tumor that has an R = 4r size would induce 5% mechanical strain on the adjacent astrocyte subunit. That 5% mechanical strain would cause the glymphatic transport within the astrocyte to drop about ∼3% (from 0.0440 μm^3^/s to 0.0428 μm^3^/s). Utilizing the same method, one can use the distribution of mechanical stress and strain in the glymphatic unit shown in Fig. 2C to estimate the glymphatic flow reduction caused by the tumor.

**Figure 5:**
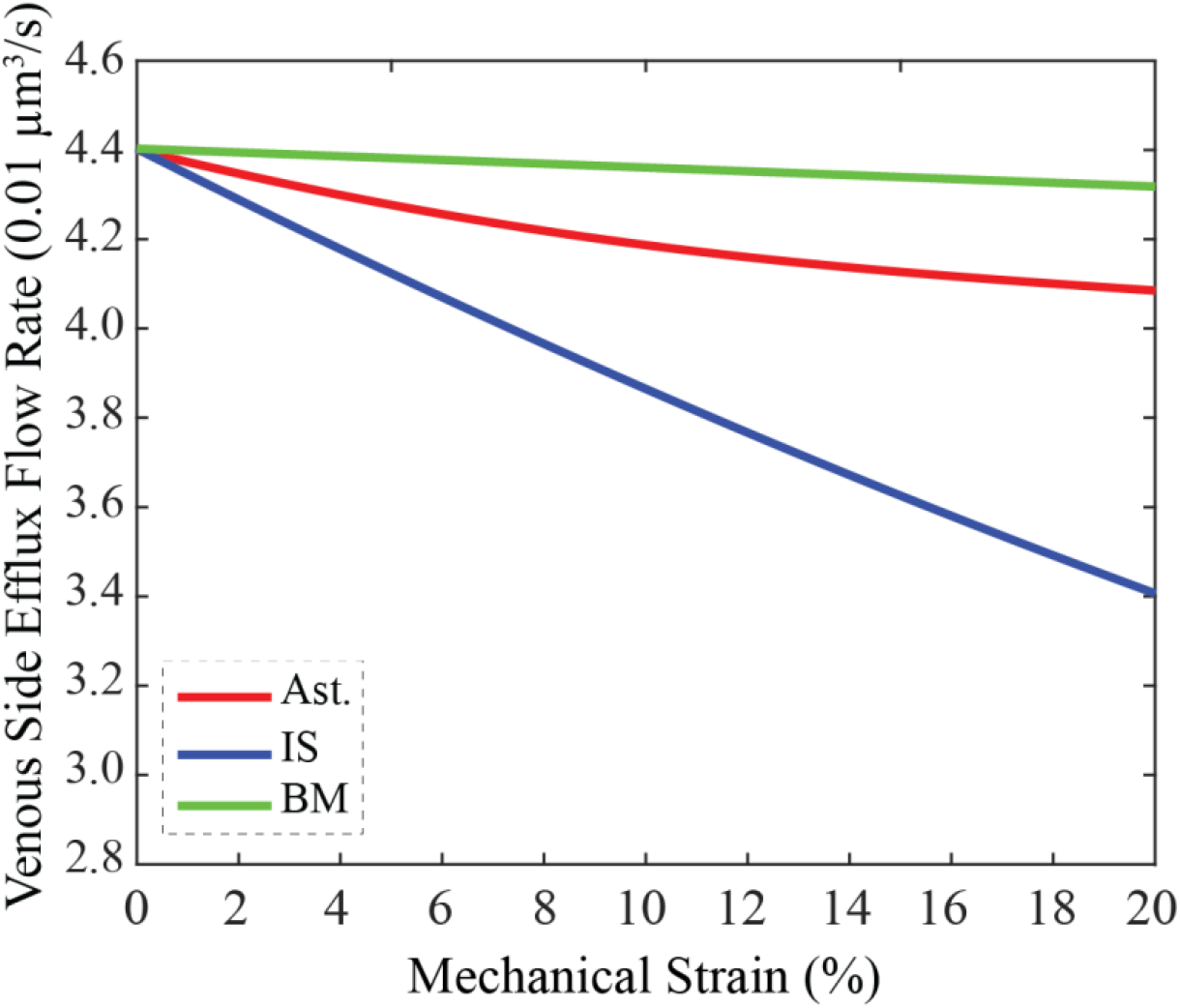
Glymphatic Efflux Flow Rate Decreases with Mechanical Strain Across Glymphatic Subunits. Venous side efflux flow rate decreases as a function of mechanical strain for the astrocyte (Ast., red), interstitial space (IS, blue), and basement membrane (BM, green) glymphatic unit components.

## Discussion

We have shown that the solid stress arising from a brain tumor acts uniquely on the different constituents of the glymphatic system and may be responsible for the experimentally observed reduction in glymphatic flow rate. This is the first model to our knowledge that incorporates the mechanical phenomena associated with brain tumors into a multicomponent fluidic model of glymphatic transport.

As a brain tumor grows, it pushes against the surrounding solid brain tissue, which is confined by a rigid skull, resulting in solid stress [11, 41]. This physical compression can directly impact the structural integrity of astrocytes, capillaries, and the interstitial spaces, disrupting their normal function within the glymphatic pathway. Compression may also impede fluid flow by altering the physical properties of the brain parenchyma, affecting the movement of substances within the glymphatic system [42, 43]. The glymphatic system’s efficiency is critical for brain health, and disruptions in waste clearance mechanisms could cause the accumulation of harmful chemical species, such as amyloid-beta and tau proteins, leading to various neurological disorders [5, 44].

As anticipated, the magnitude of stress generation increases with increasing tumor size and proximity, across all glymphatic subunits. Astrocytes emerge as critical players in maintaining the integrity of the glymphatic unit. When astrocyte contribution is minimal, stress and strain increase for the entire glymphatic unit. Increased strain in turn reduces the predicted fluid flow rate. Conversely, increased astrocyte abundance reduces mechanical strain and makes the glymphatic unit less sensitive to tumor size and proximity. Because the relative density of astrocytes varies between different regions of the brain, this may serve as a useful patient-specific indicator of glymphatic vulnerability. For example, if a patient has a tumor in or near the cerebral cortex or cerebellum, where fewer astrocytes are observed, they may be more likely to experience compression-induced glymphatic hindrance compared to a patient with a tumor in a higher astrocyte-density region such as the hypothalamus [45, 46]. In our models, we altered the astrocyte content by changing the ratio of astrocytes to IS. Thus, when astrocyte content is decreased (making the glymphatic unit more vulnerable to compression), the interstitial space is increased. This matches *in vivo* <u>observations</u>, as gliomas increase IS volume, which is also correlated with malignancy [47]. The IS does not only change in volume, however. Glioma-associated IS features overabundant ECM glycoproteins such as tenascins, as well as a more tortuous geometry, which may further hinder the movement of solutes in the interstitial fluid [47, 48]. Thus, the concurrent mechanical effects of decreasing astrocytes and the molecular alterations in the enlarged IS may synergize to reduce fluid flow and solute transport, with implications for both clearance and drug delivery.

Additionally, brain tumors can influence astrocyte behavior, resulting in both hypertrophy and hyperplasia, or conversely, astrocyte atrophy and reduced density. These changes in astrocyte morphology and distribution are often correlated with the tumor’s grade and location, contributing to the complex cellular milieu within the tumor microenvironment [49-51].

It is also important to consider the nonlocal, macroscale adverse effects of brain tumors on the CSF dynamics in the brain parenchyma [52, 53]. Brain tumors located in or near ventricles or cerebrospinal fluid pathways, can impede the normal bulk circulation of CSF. This obstruction not only affects the glymphatic function but also may lead to elevated intracranial pressure causing several side effects such as brain herniation [54], altered consciousness [55], cerebral edema [56], and hemodynamic instability [57].

A growing body of research has demonstrated the significance of solid stress in the tumor microenvironment and its impact on therapeutic efficacy [25, 58, 59]. We recently showed that losartan, an angiotensin system inhibitor normalizes the glioblastoma tumor microenvironment by decreasing ECM and solid stress levels and improving perfusion in and around the tumor [24]. Reduction of physical barriers in the ECM could enhance glymphatic system function as well. Our computational modeling approach, which incorporates mechanical stress and strain profiles applied to the glymphatic system, serves as a robust foundation for evaluating the impact of solid stress-targeting therapeutics such losartan on glymphatic function. Other applications include informing clinicians on the mechanical-glymphatic vulnerabilities of each patient or estimating brain drug delivery dynamics based on patient-specific tumor size and location information.

## Conclusions

By advancing our understanding of the mechanical underpinnings of glymphatic impairment in the context of brain tumors, this work opens up potential avenues for the development of novel therapeutic strategies to restore proper waste clearance mechanisms in the brain and improve clinical outcomes for patients with brain tumors and other neurological disorders. Clinically, this information can guide medical practitioners in anticipating complications due to hindered fluid transport, thereby informing treatment strategies and interventions. The insights gained from this parametric analysis contribute to the broader understanding of glymphatic system biomechanics, paving the way for advancements in personalized medicine and tailored therapeutic approaches for individuals suffering from brain tumors.

## Acknowledgments

We thank Ms. R’nld Rumbach for her technical support and Mr. Julian Najera for his input.

## References

[1] S. Da Mesquita, Z. Fu, and J. Kipnis, “The meningeal lymphatic system: a new player in neurophysiology,” Neuron, vol. 100, no. 2, pp. 375–388, 2018.

[2] A. Louveau, B. A. Plog, S. Antila, K. Alitalo, M. Nedergaard, and J. Kipnis, “Understanding the functions and relationships of the glymphatic system and meningeal lymphatics,” The Journal of clinical investigation, vol. 127, no. 9, pp. 3210–3219, 2017.

[3] D. Raper, A. Louveau, and J. Kipnis, “How do meningeal lymphatic vessels drain the CNS?,” Trends in neurosciences, vol. 39, no. 9, pp. 581–586, 2016.

[4] S. Da Mesquita et al., “Functional aspects of meningeal lymphatics in ageing and Alzheimer’s disease,” Nature, vol. 560, no. 7717, pp. 185–191, 2018.

[5] J. J. Iliff et al., “A Paravascular Pathway Facilitates CSF Flow Through the Brain Parenchyma and the Clearance of Interstitial Solutes, Including Amyloid β,” Science Translational Medicine, vol. 4, no. 147, pp. 147ra111–147ra111, 2012.

[6] K. Liu et al., “Attenuation of cerebral edema facilitates recovery of glymphatic system function after status epilepticus,” (in eng), JCI Insight, vol. 6, no. 17, Sep 8 2021.

[7] X. Liu et al., “Polyunsaturated fatty acid supplement alleviates depression-incident cognitive dysfunction by protecting the cerebrovascular and glymphatic systems,” Brain, Behavior, and Immunity, vol. 89, pp. 357–370, 2020/10/01/ 2020.

[8] B. T. Kress et al., “Impairment of paravascular clearance pathways in the aging brain,” Annals of Neurology, vol. 76, no. 6, pp. 845–861, 2014.

[9] H. Benveniste, X. Liu, S. Koundal, S. Sanggaard, H. Lee, and J. Wardlaw, “The glymphatic system and waste clearance with brain aging: a review,” Gerontology, vol. 65, no. 2, pp. 106–119, 2019.

[10] B. A. Plog and M. Nedergaard, “The glymphatic system in central nervous system health and disease: past, present, and future,” Annual Review of Pathology: Mechanisms of Disease, vol. 13, pp. 379–394, 2018.

[11] G. Seano et al., “Solid stress in brain tumours causes neuronal loss and neurological dysfunction and can be reversed by lithium,” (in eng), Nat Biomed Eng, vol. 3, no. 3, pp. 230–245, Mar 2019.

[12] J. Kaur et al., “Imaging glymphatic response to glioblastoma,” Cancer Imaging, vol. 23, no. 1, p. 107, 2023/10/30 2023.

[13] C. H. Toh and T. Y. Siow, “Factors Associated With Dysfunction of Glymphatic System in Patients With Glioma,” (in English), Frontiers in Oncology, Original Research vol. 11, 2021-September-23 2021.

[14] D. Xu, J. Zhou, H. Mei, H. Li, W. Sun, and H. Xu, “Impediment of Cerebrospinal Fluid Drainage Through Glymphatic System in Glioma,” (in English), Frontiers in Oncology, Original Research vol. 11, 2022-January-10 2022.

[15] P. Jin and J. M. Munson, “Fluids and flows in brain cancer and neurological disorders,” WIREs Mechanisms of Disease, vol. 15, no. 1, p. e1582, 2023.

[16] Q. Ma et al., “Lymphatic outflow of cerebrospinal fluid is reduced in glioma,” Scientific Reports, vol. 9, no. 1, p. 14815, 2019/10/15 2019.

[17] L. Li et al., “Glymphatic transport is reduced in rats with spontaneous pituitary tumor,” (in eng), Front Med (Lausanne), vol. 10, p. 1189614, 2023.

[18] C. H. Toh, T. Y. Siow, and M. Castillo, “Peritumoral Brain Edema in Meningiomas May Be Related to Glymphatic Dysfunction,” (in English), Frontiers in Neuroscience, Original Research vol. 15, 2021-April-22 2021.

[19] T. M. Mathiisen, K. P. Lehre, N. C. Danbolt, and O. P. Ottersen, “The perivascular astroglial sheath provides a complete covering of the brain microvessels: an electron microscopic 3D reconstruction,” Glia, vol. 58, no. 9, pp. 1094–1103, 2010.

[20] A. K. Shetty and G. Zanirati, “The Interstitial System of the Brain in Health and Disease,” (in eng), Aging Dis, vol. 11, no. 1, pp. 200–211, Feb 2020.

[21] N. N. Haj-Yasein et al., “Glial-conditional deletion of aquaporin-4 (<i>Aqp4</i>) reduces blood–brain water uptake and confers barrier function on perivascular astrocyte endfeet,” Proceedings of the National Academy of Sciences, vol. 108, no. 43, pp. 17815–17820, 2011.

[22] M. Asgari, D. D. Zélicourt, and V. Kurtcuoglu, “How astrocyte networks may contribute to cerebral metabolite clearance,” Scientific reports, vol. 5, no. 1, p. 15024, 2015.

[23] H. T. Nia, L. L. Munn, and R. K. Jain, “Physical traits of cancer,” Science, vol. 370, no. 6516, p. eaaz0868, 2020.

[24] M. Datta et al., “Losartan controls immune checkpoint blocker-induced edema and improves survival in glioblastoma mouse models,” Proceedings of the National Academy of Sciences, vol. 120, no. 6, p. e2219199120, 2023.

[25] H. T. Nia et al., “In vivo compression and imaging in mouse brain to measure the effects of solid stress,” Nature Protocols, pp. 1–20, 2020.

[26] L. E. Kun, “Brain tumors: challenges and directions,” Pediatric Clinics of North America, vol. 44, no. 4, pp. 907–917, 1997.

[27] L. Weninger, O. Rippel, S. Koppers, and D. Merhof, “Segmentation of brain tumors and patient survival prediction: Methods for the brats 2018 challenge,” in Brainlesion: Glioma, Multiple Sclerosis, Stroke and Traumatic Brain Injuries: 4th International Workshop, BrainLes 2018, Held in Conjunction with MICCAI 2018, Granada, Spain, September 16, 2018, Revised Selected Papers, Part II 4, 2019: Springer, pp. 3–12.

[28] L. M. DeAngelis, “Brain tumors,” New England journal of medicine, vol. 344, no. 2, pp. 114–123, 2001.

[29] J. L. Fisher, J. A. Schwartzbaum, M. Wrensch, and J. L. Wiemels, “Epidemiology of brain tumors,” Neurologic clinics, vol. 25, no. 4, pp. 867–890, 2007.

[30] P. A. Forsyth and J. B. Posner, “Headaches in patients with brain tumors: a study of 111 patients,” Neurology, vol. 43, no. 9, pp. 1678–1678, 1993.

[31] M. Wrensch, Y. Minn, T. Chew, M. Bondy, and M. S. Berger, “Epidemiology of primary brain tumors: current concepts and review of the literature,” Neuro-oncology, vol. 4, no. 4, pp. 278–299, 2002.

[32] J. M. Munson and A. C. Shieh, “Interstitial fluid flow in cancer: implications for disease progression and treatment,” (in eng), Cancer Manag Res, vol. 6, pp. 317–28, 2014.

[33] C. Pacheco, C. Martins, J. Monteiro, F. Baltazar, B. M. Costa, and B. Sarmento, “Glioblastoma Vasculature: From its Critical Role in Tumor Survival to Relevant in Vitro Modelling,” (in English), Frontiers in Drug Delivery, Review vol. 2, 2022-February-17 2022.

[34] Rakesh K. Jain, “Antiangiogenesis Strategies Revisited: From Starving Tumors to Alleviating Hypoxia,” Cancer Cell, vol. 26, no. 5, pp. 605–622, 2014.

[35] Y. Gan, J. H. Thomas, and D. H. Kelley, “Gaps in the wall of a perivascular space act as valves to produce a directed flow of cerebrospinal fluid: a hoop-stress model,” Journal of The Royal Society Interface, vol. 21, no. 213, p. 20230659, 2024.

[36] T. Koch, V. Vinje, and K.-A. Mardal, “Estimates of the permeability of extra-cellular pathways through the astrocyte endfoot sheath,” Fluids and Barriers of the CNS, vol. 20, no. 1, p. 20, 2023/03/20 2023.

[37] A. D. Martinac and L. E. Bilston, “Computational modelling of fluid and solute transport in the brain,” Biomechanics and Modeling in Mechanobiology, vol. 19, no. 3, pp. 781–800, 2020/06/01 2020.

[38] F. N. Soria, C. Miguelez, O. Peñagarikano, and J. Tønnesen, “Current Techniques for Investigating the Brain Extracellular Space,” (in English), Frontiers in Neuroscience, Review vol. 14, 2020-October-14 2020.

[39] J. M. Barnes, L. Przybyla, and V. M. Weaver, “Tissue mechanics regulate brain development, homeostasis and disease,” Journal of Cell Science, vol. 130, no. 1, pp. 71–82, 2017.

[40] K. Onwudiwe et al., “Single-cell mechanical assay unveils viscoelastic similarities in normal and neoplastic brain cells,” Biophysical Journal, 2024/03/27/ 2024.

[41] T. Stylianopoulos et al., “Causes, consequences, and remedies for growth-induced solid stress in murine and human tumors,” Proceedings of the National Academy of Sciences, vol. 109, no. 38, pp. 15101–15108, 2012.

[42] G. Lucci, A. Agosti, P. Ciarletta, and C. Giverso, “Coupling solid and fluid stresses with brain tumour growth and white matter tract deformations in a neuroimaging-informed model,” Biomechanics and Modeling in Mechanobiology, vol. 21, no. 5, pp. 1483–1509, 2022/10/01 2022.

[43] E. Comellas, S. Budday, J.-P. Pelteret, G. A. Holzapfel, and P. Steinmann, “Modeling the porous and viscous responses of human brain tissue behavior,” Computer Methods in Applied Mechanics and Engineering, vol. 369, p. 113128, 2020/09/01/ 2020.

[44] N. L. Hauglund, C. Pavan, and M. Nedergaard, “Cleaning the sleeping brain–the potential restorative function of the glymphatic system,” Current Opinion in Physiology, vol. 15, pp. 1–6, 2020.

[45] M. A. Olude, O. A. Mustapha, O. A. Aderounmu, J. O. Olopade, and A. O. Ihunwo, “Astrocyte morphology, heterogeneity, and density in the developing African giant rat (Cricetomys gambianus),” (in English), Frontiers in Neuroanatomy, Original Research vol. 9, 2015-May-26 2015.

[46] V. L. Savchenko, I. R. Nikonenko, G. G. Skibo, and J. A. McKanna, “Distribution of microglia and astrocytes in different regions of the normal adult rat brain,” Neurophysiology, vol. 29, no. 6, pp. 343–351, 1997/11/01 1997.

[47] J. Zamecnik, “The extracellular space and matrix of gliomas,” Acta Neuropathologica, vol. 110, no. 5, pp. 435–442, 2005/11/01 2005.

[48] A. S. Verkman, “Diffusion in the extracellular space in brain and tumors,” Physical Biology, vol. 10, no. 4, p. 045003, 2013/08/02 2013.

[49] P. Kleihues, F. Soylemezoglu, B. Schäuble, B. W. Scheithauer, and P. C. Burger, “Histopathology, classification, and grading of gliomas,” Glia, vol. 15, no. 3, pp. 211–221, 1995.

[50] S. C. Campbell et al., “Potassium and glutamate transport is impaired in scar-forming tumor-associated astrocytes,” Neurochemistry International, vol. 133, p. 104628, 2020/02/01/ 2020.

[51] Y. N. Diep et al., “Astrocytic scar restricting glioblastoma via glutamate-MAO-B activity in glioblastoma-microglia assembloid,” (in eng), Biomater Res, vol. 27, no. 1, p. 71, Jul 19 2023.

[52] M. Uzair-Ul-Haq et al., “Analysis of Growing Tumor on the Flow Velocity of Cerebrospinal Fluid in Human Brain Using Computational Modeling and Fluid-Structure Interaction,” arXiv preprint 2102.09742, 2021.

[53] C. F. Jones, R. S. Newell, J. H. Lee, P. A. Cripton, and B. K. Kwon, “The pressure distribution of cerebrospinal fluid responds to residual compression and decompression in an animal model of acute spinal cord injury,” Spine, vol. 37, no. 23, pp. E1422–E1431, 2012.

[54] A. Petzold and M. Smith, “High intracranial pressure, brain herniation and death in cerebral venous thrombosis,” Stroke, vol. 37, no. 2, pp. 331–331, 2006.

[55] J. Holland and R. Brown, “Acute presentation of altered conscious state,” Paediatrics and Child Health, vol. 31, no. 4, pp. 153–162, 2021/04/01/ 2021.

[56] M.-L. Ho, R. Rojas, and R. L. Eisenberg, “Cerebral Edema,” American Journal of Roentgenology, vol. 199, no. 3, pp. W258–W273, 2012.

[57] B. Pineda, C. Kosinski, N. Kim, S. Danish, and W. Craelius, “Assessing Cerebral Hemodynamic Stability After Brain Injury,” 2018, Cham: Springer International Publishing, in Intracranial Pressure & Neuromonitoring XVI, pp. 297–301.

[58] H. T. Nia, M. Datta, G. Seano, P. Huang, L. L. Munn, and R. K. Jain, “Quantifying solid stress and elastic energy from excised or in situ tumors,” Nature Protocols, vol. 13, no. 5, pp. 1091–1105, 2018/05/01 2018.

[59] H. T. Nia et al., “Solid stress and elastic energy as measures of tumour mechanopathology,” Nature Biomedical Engineering, vol. 1, no. 1, p. 0004, 2016/11/28 2016.

